# Not All Is Lost: Resilience of Microbiome Samples to Freezer Failures and Long-term Storage

**DOI:** 10.1101/2024.07.15.603584

**Authors:** M. Fabiola Pulido-Chavez, James W. J. Randolph, Sydney I. Glassman

## Abstract

Advances in technology have facilitated extensive sample collection to explore microbiomes across diverse systems, leading to a growing reliance on ultracold freezers for storing both samples and extracted DNA. However, freezer malfunctions can jeopardize data integrity. In this study, we investigated the effects of an unforeseen short-term thaw event (∼1 week) resulting from a freezer malfunction on soil samples stored at -80°C and extracted uncompromised DNA stored long-term at -20°C. We compared these samples against previously sequenced Illumina MiSeq data to assess whether the process of thawing soil or multi-year extracted DNA storage affected the resilience of bacterial or fungal richness or community composition, thereby impacting our ability to accurately determine treatment effects. Our results reveal substantial resilience in fungal richness and both bacterial and fungal beta-diversity to soil sample thawing and extended frozen DNA storage. This resilience facilitated biological inferences that closely mirrored those observed in the original study. Notably, fungi exhibited greater resilience to short-term thawing compared to bacteria, which showed sensitivity to both thawing and long-term freezing. Moreover, taxonomic composition analysis revealed the persistence of dominant microbial taxa under thawing and prolonged freezing, suggesting that dominant microbes remain viable for tracking across temporal studies. In conclusion, our study highlights that beta-diversity is more robust than alpha-diversity and fungi are more resilient than bacteria. Furthermore, our findings underscore the effectiveness of soil storage at -80°C compared to storage of extracted DNA at -20°C, despite potential freezer failures, as the most robust method for long-term storage in microbiome studies.

## Introduction

Advances in technology and affordability of molecular and ’omics’-based techniques have led to a surge in microbiome research across systems including soils (1, 2), water (3), air (4, 5), atmosphere (6), smoke (7), animals (3, 8), and humans (9, 10). Yet, sample storage techniques can affect the diversity and taxonomic composition obtained from bacterial and fungal environmental (11, 12) and human samples (Hickl et al., 2019; Plauzolles et al., 2022). Thus, microbiome research has adapted the methodology of extracting and storing microbial DNA at -20°C immediately after collection (12) or storing environmental samples at -80°C when the latter is not feasible (3), to retain the microbial composition of fresh samples (13, 14). However, the increased dependency on freezers creates an added problem with sample storage: the potential loss of stored samples due to an unexpected freezer failure, which is particularly important for long-term studies investigating microbial community compositional changes over time. Understanding how freezer storage and failure can affect soil microbial samples is necessary to ensure that ecological conclusions are appropriately inferred when working with long-term studies or with previously frozen samples or DNA.

Soil sample storage temperature (15–17) and duration (15) can affect soil microbial diversity (15), activity (18) and taxonomic composition (19). While the current body of research lacks consensus regarding which microbial groups exhibit greater resilience, and under which conditions bacteria may be more resilient than fungi and vice versa, differences in microbial physiology suggest that bacteria samples are more resilient to small changes in temperature than fungi and might be less affected by variation in soil storage techniques or temperature (Pavlovska et al., 2021; Frøslev et al., 2022). Nevertheless, while fungi and bacteria can enter a dormant state under environmental stress (20), rapid growth can occur when the environment becomes favorable, such as with increased soil moisture resulting from soil thawing (20). Thus, the response of the soil microbes to thawing can potentially obscure signals of prospective studies. However, it is worth noting that at least one study suggests that variation between treatments was greater than variation among storage conditions for soil, human fecal, and human skin samples (14), implying that an unexpected freezer failure might not significantly affect the reliability of stored microbial samples when inferring treatment effects.

The dependency on lab freezers for long-term storage of soil samples and extracted DNA for downstream analysis makes samples vulnerable to freezer failure. Given the propensity of freezer malfunctions in laboratories worldwide, it is critical to understand how short thawing events can impact microbiome samples, resulting in data loss or erroneous conclusions. Similarly, we must understand if long-term storage of frozen DNA can impact future ‘omics work. Here, we analyzed 20 soil samples that were previously sequenced using Illumina MiSeq technology targeting 16S and ITS2 regions to investigate bacterial and fungal diversity and composition (Pulido-Chavez et al., 2023). The DNA extracted from these samples had been stored for 2 years in an uncompromised -20°C freezer and excess soil from these samples was stored long-term in an -80°C ultracold freezer that experienced a complete thaw event (∼1 week) due to a freezer malfunction. This allowed us to compare the effects of short-term thawing of soil stored at -80°C vs the long-term storage of extracted DNA stored at -20°C on the alpha and beta diversity of bacteria and fungi. It also facilitated our assessment of how these storage conditions influenced our ability to draw ecological conclusions based on treatment effects and taxonomic composition.

## Methods

On November 20, 2020, we learned that our -80°C ultracold freezer that stored many of our soil samples malfunctioned, with internal temperatures of 25°C for up to 1 week, thawing all freezer contents. We took this opportunity to test how a short thawing event for unextracted soil samples stored at -80°C or multi-year storage at -20°C of extracted DNA that was not subject to freezer failure affected bacterial vs fungal resilience and our ability to determine treatment effects. We selected 20 soil samples from which DNA had previously been extracted, sequenced, and analyzed (“original” samples) to test against soil samples that had been stored in the -80°C freezer that thawed (“thawed” samples) vs resequencing extracted DNA that had been stored in an uncompromised -20°C freezer for roughly 2 years (“frozen” samples). We selected 20 previously analyzed soil samples that had been collected from burned vs unburned plots at 25, 34, 67, 95, and 286 days after the 2018 Southern California Holy Fire, selecting 2 burned and 2 unburned samples for each of the 5 timepoints (21).

### DNA extraction, sequencing, and bioinformatic and statistical analysis

In October 2021, we extracted DNA from the 20 soil samples that had experienced the short thaw, using the same methods as the “original” and “frozen” samples with Qiagen DNeasy PowerSoil Kits following the manufacturer’s protocol, as was done in the original study (21). We then used the same methods as had been used for the “original” samples to PCR amplify and sequence with Illumina MiSeq the 16S and ITS2 regions of the “thawed” and “frozen” DNA samples (21). We performed bioinformatics in QIIME2 (22) on all 60 samples (“original” vs “frozen” vs “thawed”) and performed downstream analyses on bacterial ASV tables rarefied to 6,493 sequences/sample and fungal (5,786 sequences/sample) as previously published (21).

Statistical analyses were performed in R (version 4.1.2) and all scripts can be accessed on GitHub (https://github.com/pulidofabs/Freeze-Thaw-FreezerMalfunction). We tested for correlations among bacterial vs fungal alpha diversity from the “original” vs “frozen” vs “thawed” samples using cor.test function in R with Pearson correlations (23). We further tested for correlations among bacterial vs fungal beta diversity among the treatments with mantel correlations in the Vegan package (24) using Bray-Curtis dissimilarity matrices. We then assessed how the storage types affected ecological inference based on our main treatment effects of burned vs unburned plots. For species richness, we tested treatment effects using the lme4 package (25) controlling for nestedness level of plot and time since fire. For microbial composition, treatment effects were tested using PERmutational Multivariate ANalysis of Variance (PERMANOVA) as employed by the adonis2 function in vegan (Oksanen *et al.*, 2022). We tested how the major taxonomic groups assessed in burned vs unburned plots varied in “original” vs “frozen” vs “thawed” with relative abundance plots created in ggplot2 (26) as had been done on the “original” study (21).

## Results

### Resilience of bacterial and fungal alpha diversity

In all cases, richness was strongly and significantly positively correlated to the original samples (Pearson’s R >0.6) for both bacteria and fungi (Fig. 1). Moreover, bacterial (Fig. 1a) and fungal (Fig. 1b) richness were highly resilient to soil thawing, with both taxonomic groups largely maintaining the original species richness. Overall, storing soils at -80°C regardless of freezer malfunction is more robust for fungal richness (Pearson’s R= 0.93;Fig. 1b) than storing frozen DNA at -20°C (Pearson’s R= 0.61;Fig. 1d), whereas both conditions had equivalent impacts on bacterial richness (Pearson’s R=61-64;Fig 1a,c). The correlation between frozen and thawed conditions was slightly higher for bacterial (Pearson’s R = 0.71; Fig. 1e) than fungal species richness (Pearson’s R = 0.65; Fig. 1f).

**Figure 1.**
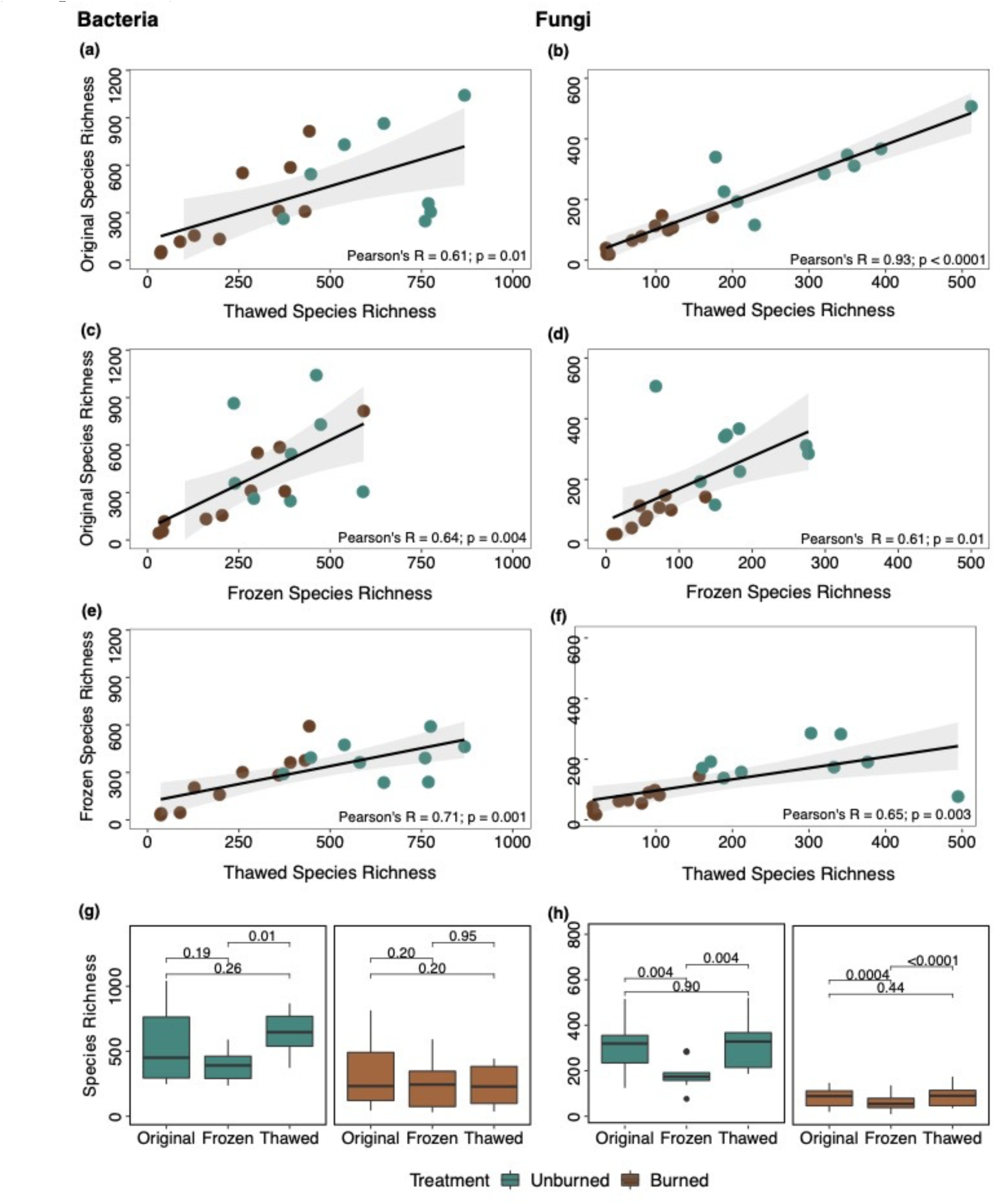
Comparison of bacterial and fungal species richness in unburned (blue-green) versus burned (brown) plots under different conditions: “original”, “frozen”, and “thawed” samples for bacteria (left panel) and fungi (right panel). Pearson correlations depict relationships between richness measures for (a) bacterial and (b) fungal “original vs thawed= samples (c) bacterial and (d) fungal “original vs frozen= and (e) bacterial and (f) fungal <frozen vs thawed” samples and (g) bacterial and (h) fungal species richness comparison between conditions, within treatments. Boxplots represent the range where the middle 50% of all values lie, with the lower end of the box representing the 1st quartile (25th percentile) and the upper end representing the 3rd quartile (75th percentile).

Even though soil stored at -80°C experienced a short thaw, soils were more resilient than storage of extracted DNA in an uncompromised -20°C freezer. Indeed, there were never significant differences among median per sample richness for original vs thawed samples for either bacteria (Fig. 1g) or fungi (Fig. 1h) for either treatment condition. In contrast, DNA that had been extracted and stored in an uncompromised -20°C freezer for up to 2 years yielded significantly lower fungal richness than original or thawed in both burned and unburned soils (Fig. 1h) and lower bacterial richness than thawed in unburned soils (Fig.1g; Table S1). Overall, original fungal richness was largely unchanged from soils that experienced a short thaw event, whereas bacterial richness was more sensitive to the thaw event.

### Resilience of bacterial and fungal beta-diversity

Beta-diversity is robust to thawing and freezing for both microbial groups, with strong positive correlations observed among original and thawed soil samples for both bacteria (R = 0.96; Fig. 2a) and fungi (R = 0.94; Fig. 2b). Bacterial beta-diversity shows strong positive correlations among original and frozen samples (R = 0.90, Fig. 2c) and frozen and thawed samples (R = 0.92; Fig. 2e). Fungal beta-diversity was also strongly positively correlated but was more sensitive than bacteria with less strong correlations among original and frozen (R = 0.77, Fig. 2d) and frozen and thawed conditions (R = 0.71, Fig. 2f). Interestingly, fungi from unburned plots subjected to long-term DNA storage at -20°C appeared to be more sensitive to changes than fungi from burned plots (Fig. 2d). Plotting the NMDS of individual samples collected at each timepoint revealed nearly indistinguishable effects of either freezing or thawing for bacteria (Fig. S1) and fungi (Fig. S2) at nearly every time point with fungal frozen samples demonstrating the most differences (Fig. S2). Therefore, while the Bray-Curtis dissimilarity of both bacteria and fungi demonstrated high resilience to storage effects, bacterial composition exhibited greater resilience compared to fungi across all storage conditions (Fig. 2).

**Figure 2.**
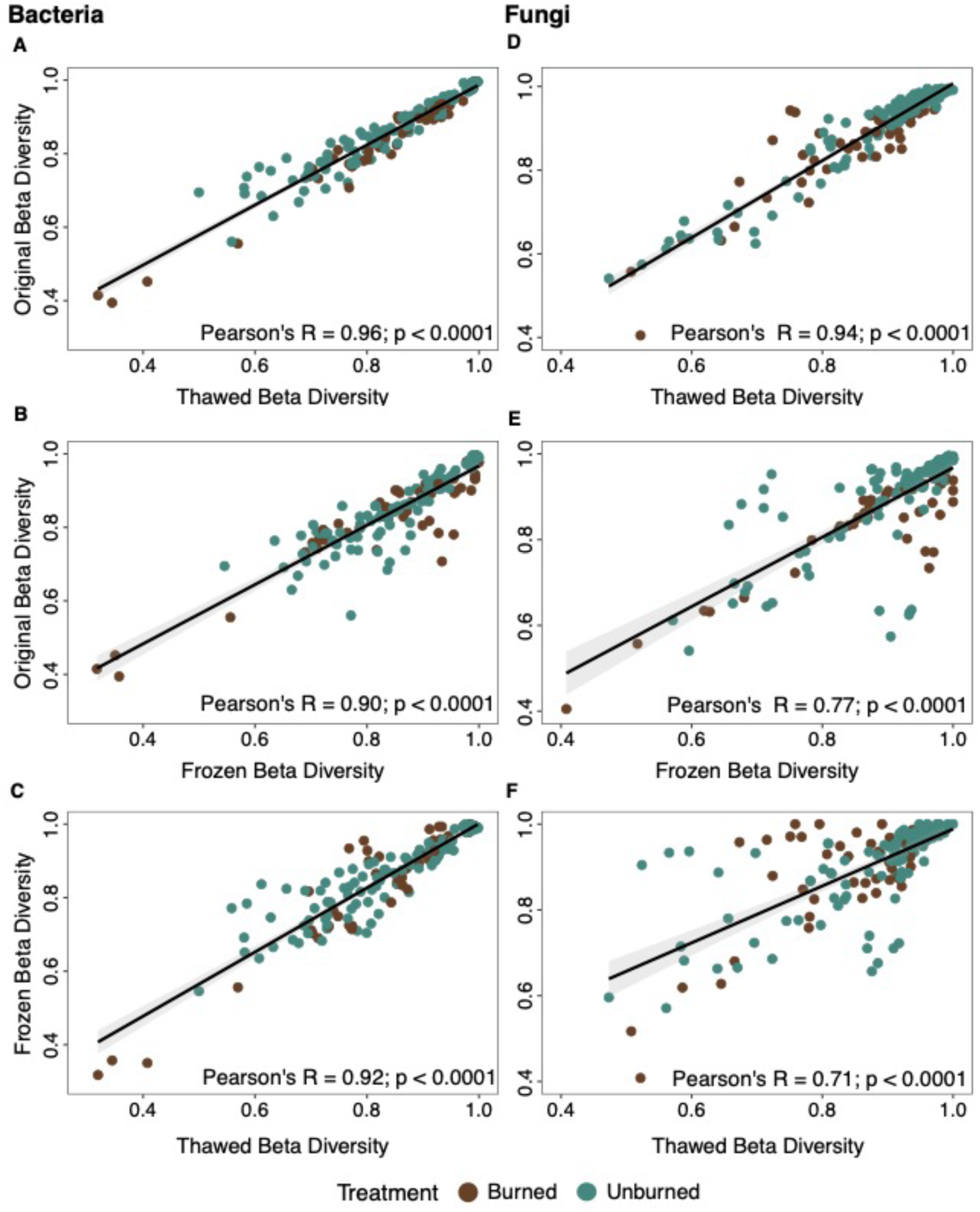
Comparison of bacterial (left panel) and fungal (right panel) beta-diversity as assessed by Bray-Curtis dissimilarity matrices. Mantel correlations compare the beta-diversity of a) bacterial and b) fungal original versus thawed, c) Bacterial and d) fungal original versus frozen and e) bacterial and d) fungal frozen versus thawed. Color denotes original treatment, with burned areas represented in brown and unburned areas in blue-green.

### Impacts of storage conditions on ecological inferences between treatment and across time

Ecological inferences, detection of treatment effects (richness averaged across time points) remained unaffected for fungi by freezing or thawing (Fig. S3). However, bacterial treatment effects were only detected in the thawed samples (Fig. S3c). For fungal richness, burned samples consistently exhibited a reduction of 66-72% compared to unburned samples regardless of storage type (Fig. S3d-e). In contrast, thawing amplified the difference in bacterial richness between burned and unburned plots, increasing the reduction from 43% (p = 0.09; Fig. S3a) in the original samples to a significant reduction of 63% in the thawed samples (p = 0.001; Fig. S3c).Treatment differences over time were also largely unaffected by storage type (Fig. 3). For bacteria, there were only significant richness reductions in burned compared to unburned samples at 44 days post-fire in all storage conditions (original, frozen, thawed), however the shapes of the curves vary slightly by storage type (Fig. 3a-c). For fungi, richness was significantly lower in burned compared to unburned samples at 3 of the 5 timepoints in original (Fig. 3d) vs 4 of the 5 timepoints for frozen (Fig. 3e) and thawed conditions (Fig. 3f). Moreover, the shapes of the curves for fungal richness are largely indistinguishable between original and thawed samples (Fig. 3d,f). However, there are slight differences observed at the first time point in frozen samples (Fig. 3e).

**Figure 3.**
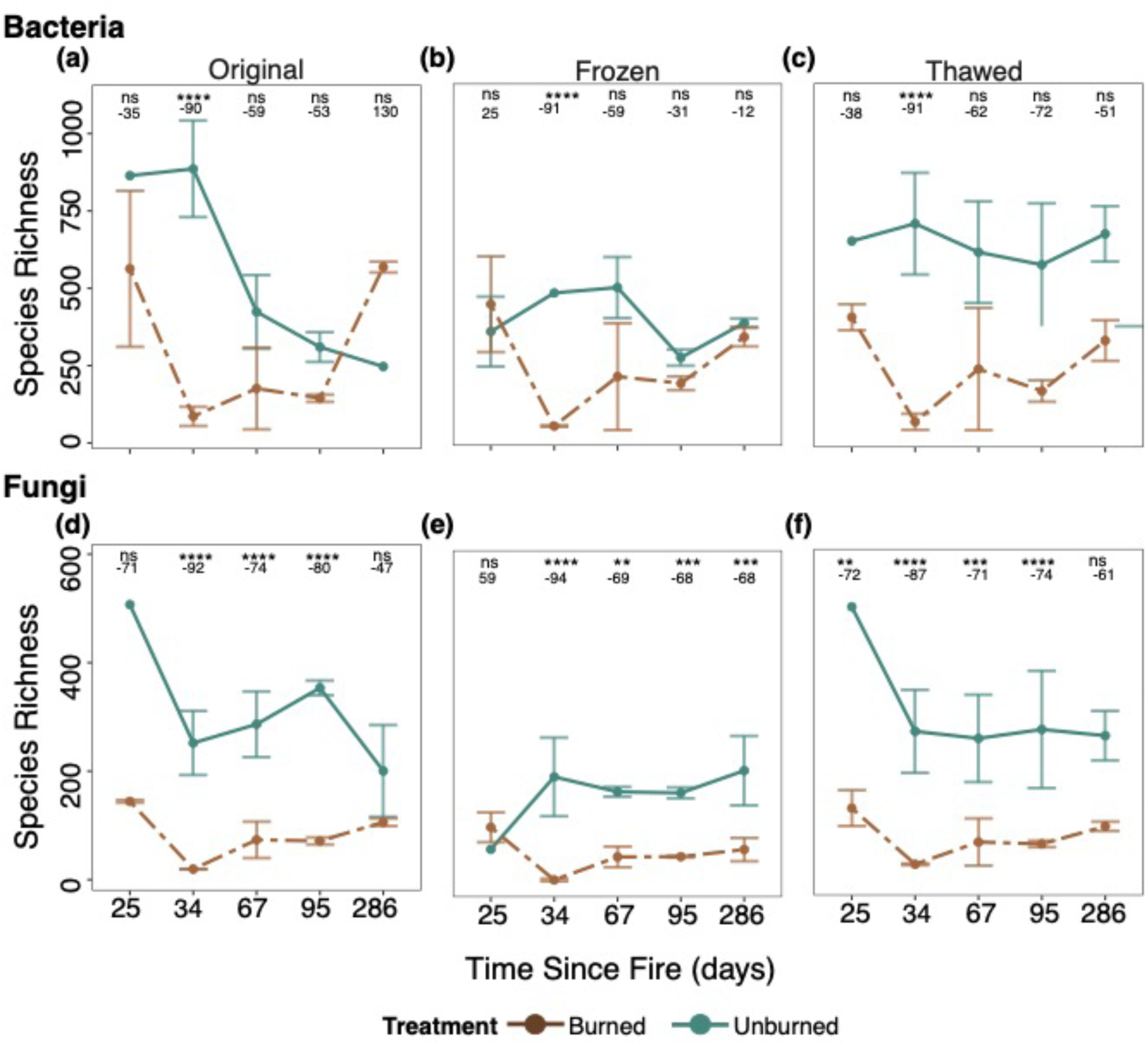
Effects of treatment (unburned in blue-green versus burned in brown) on mean bacterial and fungal species richness observed per time point for bacteria (top panels) and fungi (bottom panels) across original, frozen, and thawed storage types. Points represent mean and lines represent standard error. Significance and percent change represents difference from burned to unburned at each time point.

Differences in community composition among treatments were unaffected by storage type (Fig. 4). For bacteria, beta-diversity was always significantly different in burned vs unburned samples (p < 0.0001) with identical R^2^ values for original and thawed (R^2^=0.24) and slightly lower R^2^ in frozen (R^2^=0.20; Fig. 4a-c). Similarly for fungi, beta-diversity was always significantly different in burned vs unburned samples across all storage types (p < 0.0001) with R^2^ ranging from 0.13 to 0.19 (Fig. 4d-f).

**Figure 4:**
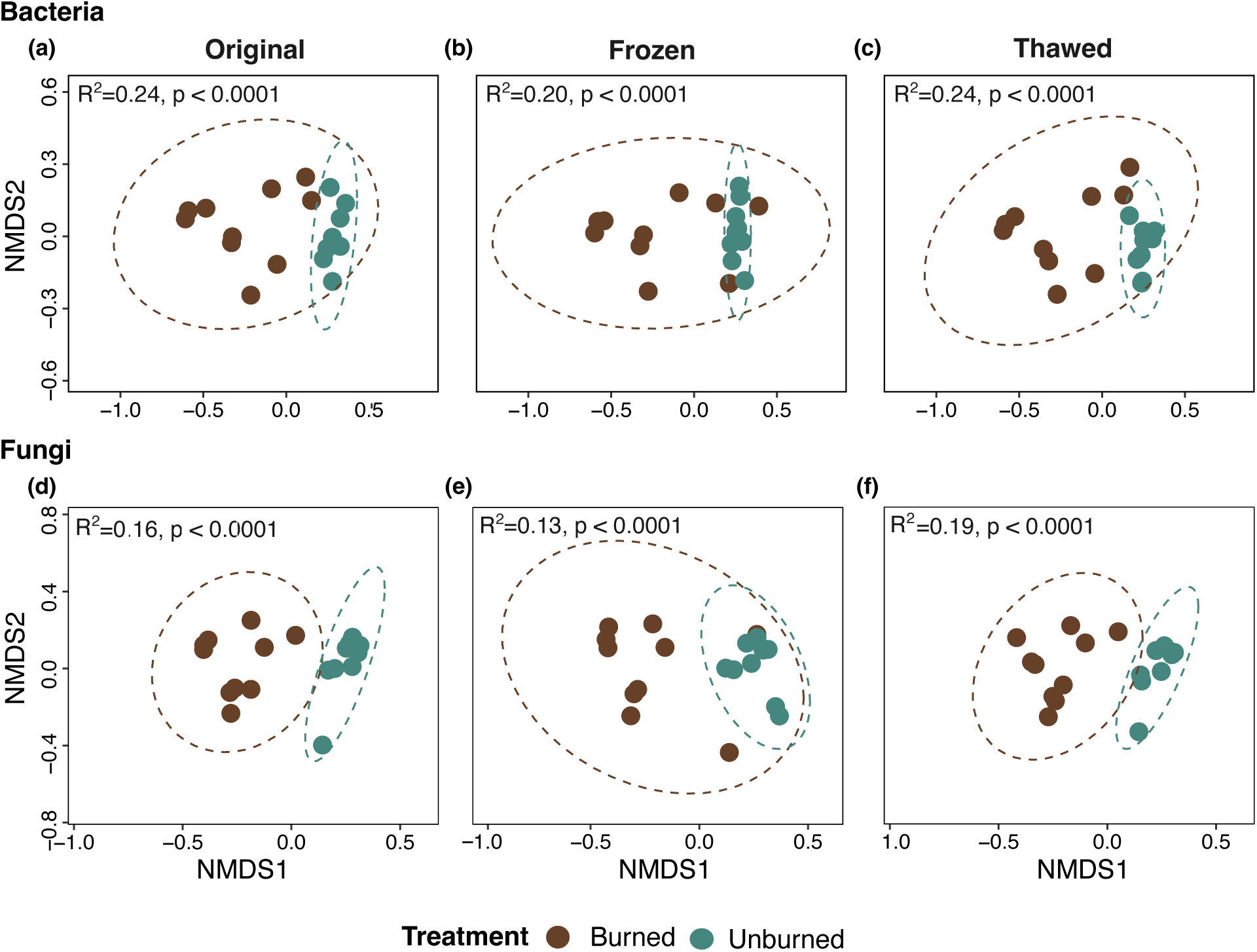
NMDS ordinations demonstrating effects of treatment (unburned in blue-green versus burned in brown) on microbial community composition across storage conditions. Impacts of fire on bacterial community composition in A) original, B) frozen, and C) thawed samples. Impacts of fire on fungal community composition in D) original, E) frozen, and F) thawed storage conditions. Ellipses represent the 95% confidence interval for each treatment based on Bray-Curtis dissimilarity. Significance based on Adonis.

### Burned communities are more resilient to change in dominance than unburned communities

Storage differences did not substantially alter the taxonomic composition of abundant genera for bacteria or fungi (Fig. 5). Similar to our findings from our original study (21), fire drastically altered the composition and dominance of bacteria and fungi with burned plots exhibiting higher dominance (Fig. 5b,d) than unburned plots (Fig. 5a,b). The taxonomic composition of genera with relative abundance above 3% (Fig. 5) and 1% (Fig. S4) from each sample type (original, frozen, or thawed) were similar for both burned and unburned communities.

**Figure 5:**
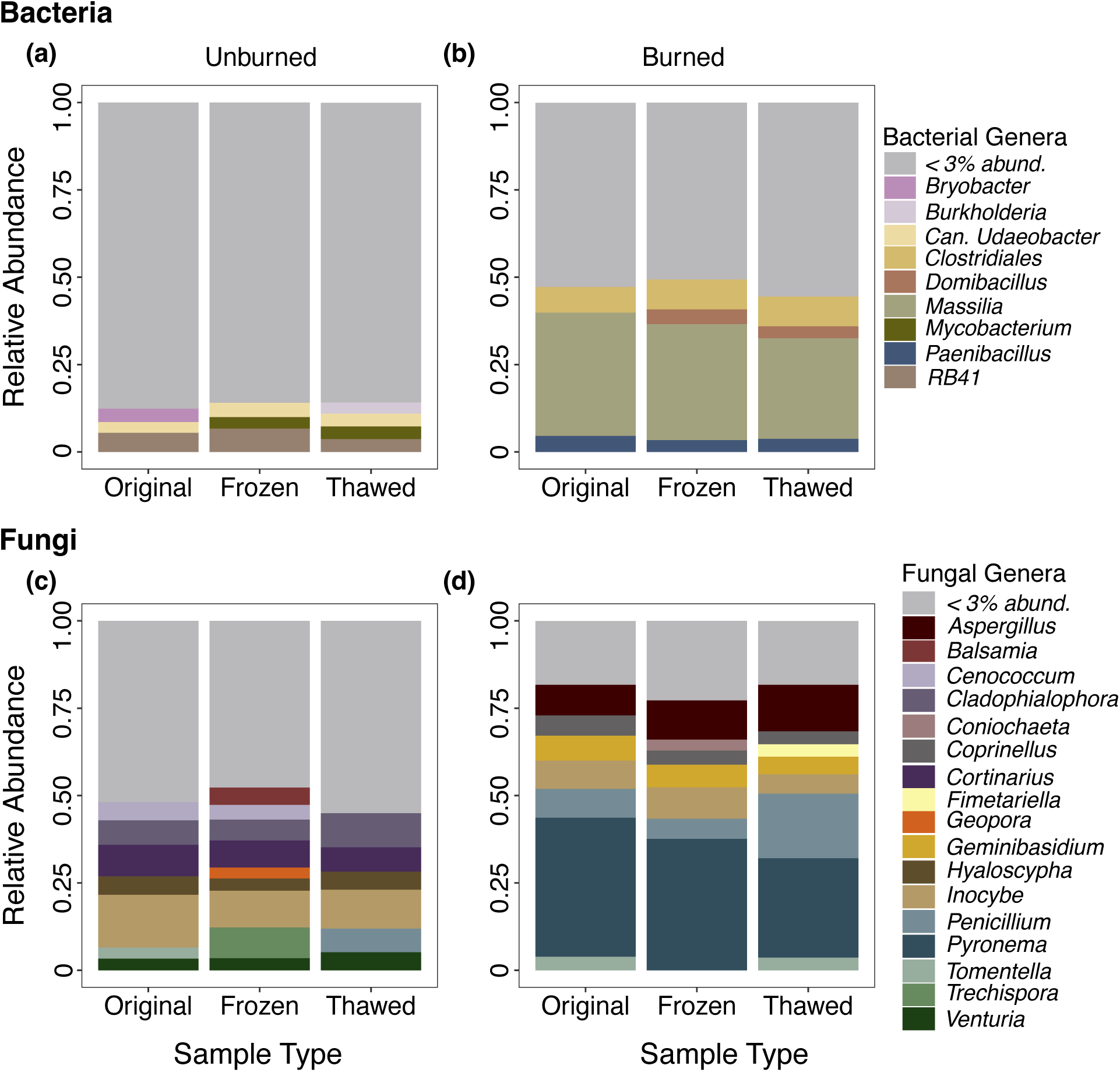
Taxonomy plots of dominant bacteria in a) unburned and b) burned plots and fungi in c) unburned and d) burned plots for original versus frozen versus thawed storage types. Genera that are greater than 3% relative sequence abundance are displayed in various colors whereas genera <3% relative sequence abundance are summarized in grey for visualization.

For bacteria, RB41 was always the most dominant bacterial genus in unburned (Fig. 5a) and *Massilia* was always the most dominant in burned plots (Fig. 5b) regardless of storage type. Comparison of all shared and unique ASVs between each sample type showed that unburned bacterial samples shared 323 ASVs (50%), while burned samples shared 48% ASVs (280; Fig. S4a, b). For fungi, *Inocybe* was always the most dominant fungal genus in unburned (Fig. 5c) and *Pyronema* was always the most dominant in burned plots (Fig. 5d) regardless of storage type. Additionally, unburned fungal samples shared 179 ASVs (56%), while burned samples shared 39% ASVs (78; Fig. S4d). While both bacterial and fungi had unique taxa for eash sample type, unique taxa were more prevalent in thawed unburned samples and frozen burned fungal samples (Fig, S4c,d).

## Discussion

Microbiome research relies heavily on freezer reliability for sample preservation, yet technological failures can jeopardize valuable datasets. In this study, we demonstrate that not all is lost when an unforeseen freezer failure occurs. Notably, correlations remained strongly and significantly positive for bacteria and fungi for both alpha (r>0.6) and beta-diversity (r>0.71) across all storage conditions. Bacterial richness was slightly more sensitive to the thaw event (r=0.61) than fungal richness (r=0.93) or bacterial (R=0.94) or fungal (R=0.96) composition, but even so ecological inferences based on treatment effects remained largely unchanged. We conclude that long-term storage of soil at -80°C is more robust than storage of extracted DNA at-20°C, even in the face of temporary ultracold freezer failure, since storing DNA at -20°C for 2 years led to decreased richness in both bacteria and fungi.

### Robustness of Ecological Inferences Despite Soil Sample Thawing

Overall, the fact that bacterial and fungal alpha metrics between the original and thawed samples were highly correlated suggest that ecological patterns and conclusions can be adequately reached. Indeed, we found consistent treatment effects on fungal species richness and community composition between the original and thawed samples, including similar decreases in richness between burned and unburned samples and alterations in community composition. Bacterial richness was more sensitive to thawing than fungal richness, which is corroborated by prior research on soils in a Chinese temperate forest (27). In our case, thawing the samples amplified treatment effects when averaging across timepoints (Fig. S3) although treatment effects were not detected at most timepoints regardless of storage type when analyzed individually, likely because we only analyzed a subset of our original samples for the purposes of this analysis (Fig. 3a). We speculate that thawing the soil magnified treatment effects because our soils came from chaparral shrublands that are adapted to arid environments (28, 29) and likely have evolved mechanisms for rapid response to increased soil moisture, such as those that occur during the thawing of soil, and thus explaining the increase in richness in unburned plots observed here. Indeed, prior research indicates that bacterial communities rapidly react to rain events (30).

### Long-Term Freezing Negatively Impacts Microbial Richness

Despite alterations in total species richness, ecological inferences based on our treatment effect remained the same across storage conditions for both bacteria and fungi (Fig. 3). However, our research indicates that correlations between original and frozen richness were lower for both bacteria (r=0.64) and fungi (r=0.61), indicating that freezing soil at -80°C is more robust than freezing extracted DNA at -20°C for long-term storage, even if the -80°C completely thawed for up to a week. Freezing DNA at -20°C is a standard practice in microbiome work (3). Although prior research suggests that freezing DNA for up to 14 days has minimal impact on the community (14), extended freezing periods are often necessary for microbial studies. Our study demonstrated that long-term freezing of extracted DNA, spanning over 2 years, resulted in decreased bacterial and fungal richness in burned and unburned samples accordingly, aligning with findings from previous short-term freezing studies (15, 19, 31). For viruses as well, freezing at -20°C compromised DNA integrity bacteriophage lambda with increased freezing time < 1 year (32). Additionally, studies suggest that physical shearing due to buffer type, influenced by pH and salt concentration, as well as freeze-thaw cycles (accessing DNA over time), can result in DNA degradation (33). As a result, for long-term temporal analysis, storing soil samples at - 80°C and extracting DNA as needed is more robust than storing extracted DNA at -20°C for extended periods.

### Neither a Short-term Thaw nor Long-Term Freezing affect Microbial Community Composition

The analysis of beta-diversity, crucial for understanding microbial community composition and treatment effects, remained highly robust across all sample storage methods. Our findings indicate that both long-term storage of extracted DNA in an uncompromised -20°C and re-extraction of soil stored at -80°C, even in the case of a failure and complete thaw, produced highly correlated beta-diversity metrics and yielded similar treatment effects to our original samples (21). This is consistent with a global meta-analysis study of soil freeze–thaw cycles of snow cover in various ecosystems (34).

Our study reveals the robustness of bacterial and fungal beta-diversity patterns across all sample types, with bacteria exhibiting greater resilience than fungi. Notably, the correlations between frozen and thawed fungal communities, and between original and frozen fungal communities, were lower compared to those observed for bacterial communities. This disparity may be attributed to the larger average base pair size of fungal DNA as compared to bacterial DNA, especially since smaller DNA sizes are known to confer improved stability against long-term freezing and increased freeze-thaw cycles (33). Despite lower correlations in fungal beta-diversity metrics, the ecological inferences of treatment effects based on community compositional differences across treatments remained the same. This suggests that researchers can confidently conduct long-term temporal studies on microbial community composition even if their soil samples experience a short-term thaw or if the extracted DNA has been stored at -20°C for long periods.

### Burned samples more robust than unburned samples to freezing and thawing

We found particularly high robustness to storage conditions for our burned samples, building upon our original study that highlighted the lower diversity and heightened stability of burned microbial communities (21). Moreover, our findings support the notion that less diverse communities are inherently more stable due to reduced stochasticity (35). Importantly, our results suggest that soil samples with initially low biological diversity retain significant ecological information even after unforeseen thawing events, underscoring their enduring value and relevance. Additionally, the presence of stress-tolerant taxa within burned microbial communities (21, 36), could further confer resilience against rapid stressors such as unexpected thaw events. This resilience highlights the adaptability of microbial communities and emphasizes the importance of considering their potential responses to environmental perturbations in ecological research. Although minor changes were observed in less dominant bacterial and fungal taxa, these changes were based on unique taxa in each sample type, which on average constituted less than 11% of the bacterial community and less than 13% of the fungal community, regardless of treatment (burned or unburned; Fig. S4). However, the distribution of dominant genera remained approximately equal across sample types, consistently making up more than 50% of shared taxa for bacteria and 48% for fungi. Thus, given that dominance is a key descriptor in burned and disturbed environments, we can confidently say that biological inferences can indeed be made using thawed and frozen samples. Furthermore, these results align with studies indicating that dominant taxa are more resistant and resilient to environmental changes. (37, 38). However, it is also possible that our particular soil samples might be more likely to be highly resistant to freezer failures. For example, the most dominant bacterial taxa in our burned samples, *Massilia*, is highly tolerant of abiotic stressors (39). The dominant fungal genera in our burned plots such as *Pyronema* produce resistant sclerotia (40) and fungal genera like *Penicillium* and *Aspergillus* are capable of producing resistant conidia (41), which may make them more likely to survive freezer failure than other sample types.

## Conclusions

Here, we compared the ecological implications of an unexpected -80°C freezer failure (∼1 week) and long-term DNA storage at -20°C (2 years) on soil microbiome studies and our ability to properly assess community dynamics and treatment effects for long-term monitoring studies. Our findings revealed high resilience of fungal richness and both bacterial and fungal beta-diversity to thawing of soil samples and long-term frozen DNA storage. The robust resilience of fungal richness facilitated biological inferences regarding treatment effects, regardless of sample type. However, for bacteria, although the patterns resembled those of the original samples, they appeared more pronounced in both frozen and thawed samples. In contrast, treatment effects were consistently detected for bacterial and fungal beta-diversity, suggesting that community composition is more resilient to small changes in environmental conditions. In conclusion, we suggest that, even in case of a short-term freezer failure, stored soil samples retain their usability for subsequent DNA based analysis, signifying promising potential for salvaging data in such situations.

## Acknowledgments

We thank the Cleveland National Forest and the Trabuco Ranger District, particularly District Ranger Darrel Vance and Emily Fudge, Jeffrey Heys, Lauren Quon, Jacob Rodriguez, and Victoria Stempniewicz, for their guidance, assistance with permitting, and support in site selection. We thank Judy A. Chung for her help with fieldwork and molecular work and Aral C. Greene, Dylan Enright, Sameer S. Saroa, and Elizah Stephens for fieldwork assistance. We thank the labs of Jennifer Martiny and Alejandra Rodriguez-Verdugo and our anonymous reviewers for their feedback on this manuscript. This work was supported by USDA NIFA grant #2022-67014-36675 to SIG.

## Conflict of interest

The authors declare no conflicts of interest.

